# Time-resolved dual RNA-Seq reveals extensive rewiring of lung epithelial and pneumococcal transcriptomes during early infection

**DOI:** 10.1101/048959

**Authors:** Rieza Aprianto, Jelle Slager, Siger Holsappel, Jan-Willem Veening

## Abstract

*Streptococcus pneumoniae* (pneumococcus) is the main etiological agent of pneumonia. Pneumococcal pneumonia is initiated by bacterial adherence to lung epithelial cells. Infection to the epithelium is a disruptive interspecies interaction involving numerous transcription-mediated processes. Revealing transcriptional changes may provide valuable insights into pneumococcal disease. Dual RNA-Seq allows simultaneous monitoring of the transcriptomes of both host and pathogen. Here, we developed a time-resolved infection model of human lung alveolar epithelial cells by *S. pneumoniae* and assessed transcriptome changes by dual RNA-Seq. Our data provide new insights into host-microbe interactions and show that the epithelial glutathione-detoxification pathway is activated by bacterial presence. We observed that adherent pneumococci, not free-floating bacteria, access host-associated carbohydrates and repress innate immune responses. In conclusion, we provide a dynamic dual-transcriptomics overview of early pneumococcal infection with easy online access (http://dualrnaseq.molgenrug.nl). Further database exploration may expand our understanding of epithelial-pneumococcal interaction, leading to novel antimicrobial strategies.

**Graphical Abstract:** 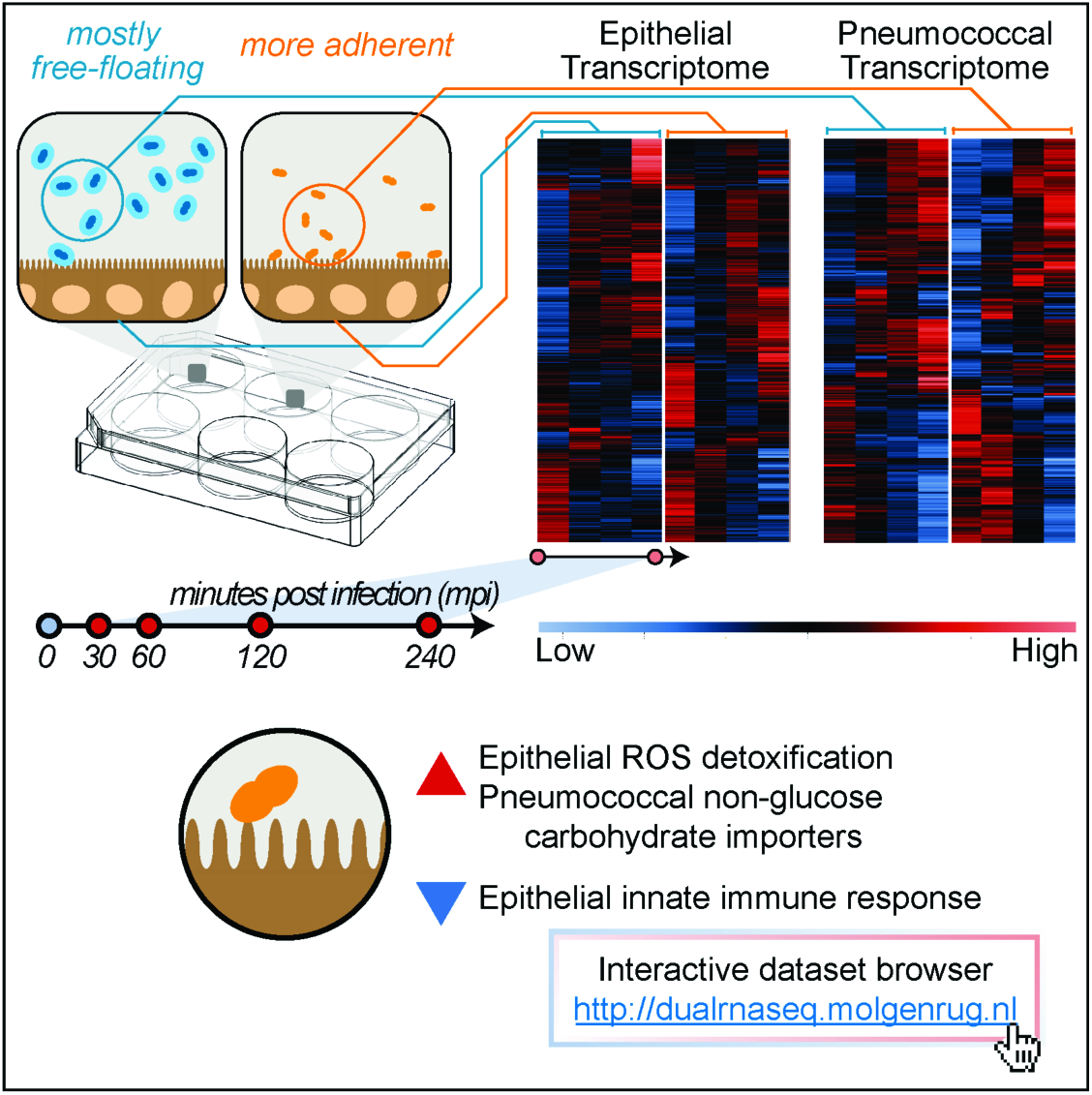

## Introduction

Lower respiratory tract infections (LRTIs), or pneumonia, claim more lives than any other communicable disease worldwide; the main etiologic agent behind this infection is the Gram-positive opportunistic pathogen *Streptococcus pneumoniae* (pneumococcus, Prina et al., 2015). Normally part of the human nasopharyngeal microflora, *S. pneumoniae* can invade the lower airways where it provokes host inflammatory and immune responses(Kadioglu et al., 2008). At the earliest stage of infection, pneumococcus adheres to epithelial cells and interacts intimately with the epithelium (Hammerschmidt et al., 2007). Meanwhile, host and pathogen cross-communicate and, they simultaneously, affect each other in a disruptive manner (Kadioglu et al., 2008; Lee et al., 2004). This interspecies interaction activates numerous processes in epithelial and pneumococcal cells (Bootsma et al., 2007; Mlacha et al., 2013). To obtain comprehensive and meaningful biological knowledge of the infection processes involved in pathogenesis, simultaneous monitoring of the transcriptome changes in both species is required (Westermann et al., 2012).

Lung epithelial cells perform vital roles during infection. First, the cells form a physical barrier to the external environment. On top of these cells, a thick layer of epithelial-derived mucus offers extra protection that traps and removes pathogens (Voynow and Rubin, 2009). Mucins, the main component of mucus, are large glycoprotein-polymers rich in sialic acids and other aminosaccharides (Rose and Voynow, 2006). Additionally, epithelial cells kill pathogens directly by producing antimicrobial peptides, e.g.: defensins and cathelicidins (Tecle et al., 2010). Moreover, epithelial cells regulate innate immune responses by secreting a wide array of pro-inflammatory cytokines, that recruit neutrophils and activate macrophages (Hallstrand et al., 2014). Finally, epithelial cells activate adaptive immune cells, including dendritic cells and T-cells via chemokine expression (Soumelis et al., 2002).

Pneumococcal adherence to epithelial cells is the first necessary step to pathogenesis (Bogaert et al., 2004). In order to adhere, pneumococcus must quickly shed the thick exopolysaccharide capsule, which protects against phagocytes (Abeyta et al., 2003; Hyams et al., 2010). The shedding exposes surface adhesion factors and desensitizes the bacterium from antimicrobial peptides (Beiter et al., 2008; Kietzman et al., 2016). Subsequently, *S. pneumoniae* must acquire nutrients to support growth and, at the same time, evade host immune responses (Shelburne et al., 2008). Pneumococcal factors may be involved in multiple processes, e.g.: PsaA, a surface-exposed protein, acts concurrently as adhesion factor and manganese transporter (Rajam et al., 2008). The scarce manganese (Gray et al., 2010) helps in neutralizing reactive oxygen species (ROS) and bacterial fitness (Tseng et al., 2002).

Interspecies interaction during infection is a chaotic process which necessitates rapid and massive adaptation for epithelial and pneumococcal survival. During the adaptation, transcriptional changes plays a focal point both in host(Jenner and Young, 2005) and in pathogen (Sorek and Cossart, 2010). RNA-sequencing (RNA-Seq) delivers genome-wide quantitative snapshots of the transcriptome (Kukurba and Montgomery, 2015). In a thought experiment, Vogel and co-workers argued that simultaneous profiling of host and pathogen transcriptomics by dual RNA-Seq might provide valuable insights for infection biology (Westermann et al., 2012). Recent dual RNA-Seq studies were successful in elucidating the role of sRNAs in the intracellular pathogen *Salmonella typhimurium* (Westermann et al., 2016), cross-talk in the Gram-negative LRTI pathogen *Haemophilus influenzae* (Baddal et al., 2015) and transcription profiles in the protozoan *Leishmania major* (Dillon et al., 2015) during their respective infection.

Here, we exploited the dual RNA-Seq approach to simultaneously monitor the transcriptome cross-talk between lung alveolar epithelial cells and pneumococci during early infection. Due to the transient and highly-dynamic nature of the transcriptome (Pedersen et al., 2011), we monitored the transcriptional changes in a time-resolved manner. Moreover, since pneumococcal adherence to epithelial cells determines the outcome of early infection, we compared adherent and non-adherent *S. pneumoniae* to allow specific transcriptional interrogation on adherence. Additionally, we confirmed our dual RNA-Seq gene expression data by qRT-PCR and quantitative fluorescence microscopy to visualize pneumococcal proteins and thereby confirm several novel biological observations identified in the dataset. Finally, we developed a user-friendly online database (http://dualrnaseq.molgenrug.nl), giving access of our detailed time-resolved dual transcriptomes data to the pneumococcal, microbiology, immunology and pulmonology research communities.

## Results

### The early pneumococcal infection model to epithelial cells

At the first phase of LRTI, *S. pneumoniae* adheres to the sterile apical-side of epithelial cells, adaptation of bacterial transcriptome occurs followed by bacterial outgrowth (Mlacha et al., 2013), which in turn stimulate epithelial transcriptional responses to their presence (Bootsma et al., 2007). We aimed to recapitulate these events in an *in vitro* model consisting of co-incubation of the pathogenic *S. pneumoniae* strain D39 (serotype 2) to a confluent layer of type II lung alveolar human epithelial cell (A549) at a multiplicity of infection (MOI) of 10, i.e., ten pneumococci per epithelial cell (**Figure 1A**). Five time points up to 4 hours after infection were selected to capture both early transcriptome responses (30 and 60 mpi) and later responses (120 and 240 mpi). Six technical replicates (individual wells) were pooled into one biological replicate. Two biological replicates were used for each time point, except for 240 mpi where we only obtained one replicate (**Figure 1E**). In order to elucidate adherence-specific expression, we incorporated an isogenic unencapsulated D39 strain (Δ*cps2E*) with increased adherence to epithelial cells into the model(Kjos et al., 2015). The capsular mutant showed significantly greater capacity to adhere to epithelial cells than its encapsulated parental strain (*p*<0.001, **Figure 1C**). During infection, the total number of cells of both strains were significantly (*p*<0.01) increased after 4 hours (**Figure 1D**), showing that both strains multiply in the model, thereby recapitulating one of the characteristics of infection. To minimize transcriptional changes because of sample handling, we did not separate cellular mixtures before total RNA isolation (epithelial cells, adherent pneumococci and free-floating pneumococci, **Figure 1E**, see **Supplemental Information**).

**Figure 1.**
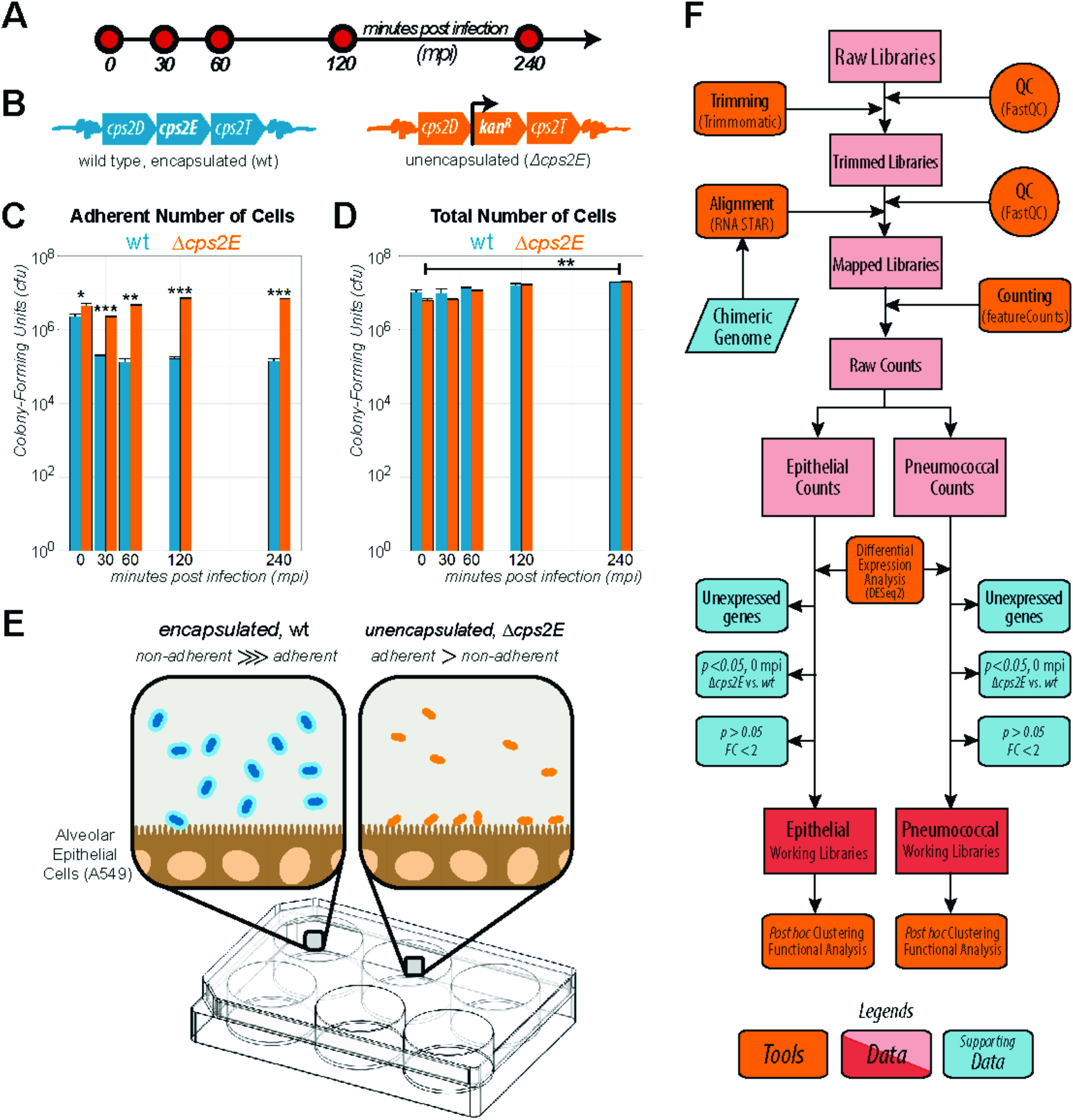
The early infection model. Confluent monolayer of alveolar epithelial cell line (A549) was co-incubated with *S. pneumoniae* strain D39. **A**. We chose five infection time points: 0, 30, 60, 120 and 240 minutes post infection (mpi). **B**. Since adherence is a hallmark of infection, we used an unencapsulated strain, created by disrupting *cps2E* vital in capsule biogenesis. **C**. Δ*cps2E* strain adhered more readily to epithelial cells than its encapsulated parental strain. 30 mpi, Δ*cps2E* (orange bar) showed significantly (*p<0.001*) more adherent cells than its parental strain (cyan bar). Data is presented as mean±SEM. **D**. At 240 mpi, both strains multiplied significantly (*p*<0.01) with no significant difference between strains. **E**. Encapsulated strain has more free-floating than adherent cells while Δ*cps2E* has a higher fraction of adherent bacteria. **F**. After quality-check, low-quality reads were trimmed. Reads were aligned to a synthetic chimeric genome. Aligned reads were counted and classified as epithelial or as pneumococcal counts. We removed three gene fractions, clustered and performed functional enrichment to the working libraries.

To analyze the time-resolved dual RNA-Seq dataset, a combination of freely available bioinformatics tools was used (**Figure 1F**). First, raw reads were trimmed (Bolger et al., 2014) and aligned (Dobin et al., 2013) to a chimeric genome containing the concatenated genome of *Homo sapiens* (GRCh38, Ensembl) and *S. pneumoniae* D39 (NC_008533.1, NCBI). The one-step mapping was chosen to minimize rates of false negatives. Reads were then separately counted (Liao et al., 2014) and classified as either epithelial or pneumococcal. Following differential gene expression analysis, three groups of genes were removed (see below) and unbiased automatic clustering (Kumar and E. Futschik, 2007) and functional enrichment were performed (Dennis et al., 2003; van Opijnen and Camilli, 2012).

### Dual RNA-Seq generates high-quality datasets with clusters of epithelial and pneumococcal co-expressed genes

Dual RNA-seq generated single end 75 nt reads. We sequenced to such depth, that on average each library has 70 million reads (30 to 95 million). After adapter trimming and removal of low-quality reads, we retained 92% (89.0-93.0%). Subsequently, 79% of reads (74.6 to 83.0%) successfully aligned to the chimeric genome. Additionally, we concluded that dual rRNA depletion was successful since only 0.03% of human and 0.24% of pneumococcal reads mapped as ribosomal RNAs. For each library, we counted 18 million epithelial reads (26%) and 37 million pneumococcal reads (52%, **Figure 2A**). The high number of reads in each library and the high fraction of usable reads highlights the quality and suitability of our approach for dual RNA-Seq: simultaneous total RNA isolation, dual rRNA depletion and cDNA library preparation. PCA analysis showed no evidence for batch effects (**Supplemental Figure 1**). Relative enrichment of pneumococcal reads may stem from the total RNA isolation protocol. Nevertheless, each library contained sufficient epithelial reads for differential gene expression analysis (The ENCODE Consortium, 2011).

**Figure 2.**
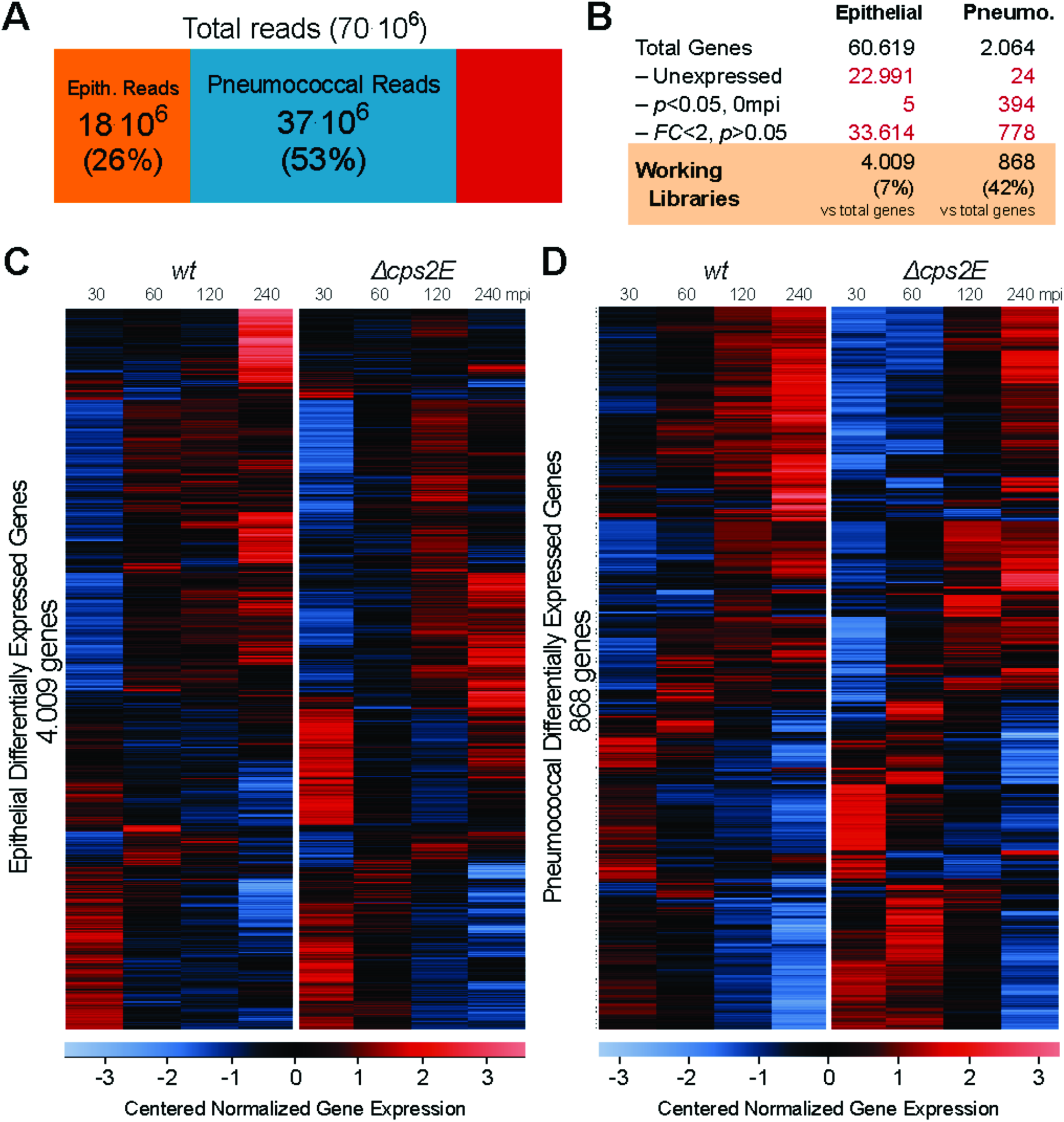
Dual RNA-Seq generates high-quality datasets suitable for probing host-pathogen transcriptomes. **A.** On average, there are 70 million reads per library: 18 million reads were mapped to the human genome while 37 million reads to pneumococcal genome. **B.** We excluded three gene fractions: unexpressed genes, i.e.: without counts in any libraries; genes that were differentially expressed at 0 mpi (*p*<0.05) between Δ*cps2E* and wt libraries and genes with no statistical significance (*p*>0.05) and fold change less than two (*FC*<2) in all contrast. Epithelial working libraries contain 4.009 genes (7% of human genes) while pneumococcal working libraries 868 genes (42% pneumococcal genes). **C.** Gene expression in epithelial working libraries were normalized, centered and automatically clustered. The left panel shows epithelial genes in response to encapsulated strain while the right panel shows epithelial response to Δ*cps2E S. pneumoniae* at different time points. Clear clusters of co-expressed epithelial genes can be observed in the heat map. Blue indicates a relatively lower expression while red, a higher value. **D**. Correspondingly, we presented pneumococcal expression in the same manner: left panel shows wild type pneumococcal response to epithelial cells, while right panel shows Δ*cps2E* response.

To simplify further analyses, we excluded three gene fractions (**Figure 2B**). First, we removed unexpressed genes, i.e., without any counts in all libraries: 22.991 (38%) epithelial genes and 24 (1%) pneumococcal genes. The relatively large fraction of unexpressed epithelial genes is in accordance with recent studies on the human epithelial transcriptome (Hackett et al., 2012; St-Pierre et al., 2013). Second, we excluded genes that were differentially expressed (*p*<0.05) at 0 mpi between unencapsulated (Δ*cps2E*) and encapsulated (wt) libraries. While only five epithelial genes were removed, 394 (19%) pneumococcal genes were already differentially expressed at 0 mpi. Although polar effect due to *cps2E* disruption can explain differential expression of genes in the 17kb long *cps* operon, it remains unknown why other genes were differentially expressed. We speculate that constructing the thick exopolysaccharide capsule requires specific transcriptional fine-tuning of numerous genes outside the *cps* locus. Finally, we removed genes with no significant difference (*p*>0.05) and genes with fold changes less than 2 in all contrasts (**Supplemental Figure 2**). In total, the epithelial working libraries contained 4.009 (7% of total) genes and pneumococcal working libraries 868 (42% of total) genes.

To compare gene expression, we normalized expression values using DESeq2 (Love et al., 2014), centered and clustered the values (Kumar and E. Futschik, 2007). The centered normalized values were visualized as heat maps, divided into two panels, one for each bacterial strain. Strikingly, heat maps showed obvious clusters of co-expressed genes and clear gene expression differences between adhering (Δ*cps2E*) and non-adhering (wt) bacteria to epithelial cells (**Figure 2CD**). Specifically, the left panel of **Figure 2C** shows the epithelial transcriptional response exposed to encapsulated *S. pneumoniae* at different time points (30, 60, 120 and 240 mpi) while the right panel displays the response in contact with the unencapsulated strain. *Vice versa*, co-expressed clusters in pneumococcal genes are differentially expressed in contact with human epithelial cells (**Figure 2D**).

Making raw data publicly-available has been common practice in recent years, as we have done for this project (GEO accession number GSE79595). Unfortunately, publicly available datasets do not translate to direct exploration and extraction of biological insights for the majority of researchers. Therefore, we built an easily-accessible online platform which hosts the complete dual RNA-Seq database. To access and visualize the data, users can simply select the gene of interest (or multiple genes of interests) and examine their expression during early infection (**Supplemental Figure 3**). Expression data can be downloaded and opened in common spreadsheet software e.g.: Microsoft Excel^®^. To visualize expression, users can choose from three normalization methods: DESeq2 normalization(Love et al., 2014), TPM (transcript per million (Wagner et al., 2012) and log_2_-transformed TPM values.

### Validation of dual RNA-Seq by qRT-PCR and pneumococcal protein fusions

To confirm dual RNA-Seq data by quantitative real-time PCR (qRT-PCR), we chose 10 epithelial genes (*ABCC2, AKR1B10, AKR1C3, ALDH1A1, DEFB1, DKK1, IDH1, NOLC1, PTGES* and *TXNRD1*) and 19 pneumococcal genes (*amiC, blpY, dinF, hrcA, infC, lytA, malC, manL, msmR, nrdD, pulA*, SPD_0249, SPD_0392, SPD_0475, SPD_0961, SPD_0990, SPD_1517, SPD_1711 and SPD_1798). The target genes were selected because of their varied expression profiles: increasing, decreasing or unchanged. The cycle threshold (Ct) for epithelial transcripts were normalized against *ACTB* (β-actin) while pneumococcal transcripts were normalized to *gyrA* (gyrase A). The reference genes were highly expressed and did not show significant changes (*p*>0.05, *FC*<2) between any time points during early infection. The qRT-PCR fold change was calculated by the ΔΔCt method (Livak and Schmittgen, 2001) to one time point. Fold changes obtained by qRT-pCR and dual RNA-Seq showed a relatively high correlation for both species: epithelial transcripts, (*R*^*2*^=0.72) and pneumococcal transcripts, (*R*^*2*^=0.73, **Figure 3A**), validating the reliability of dual RNA-Seq data.

**Figure 3.**
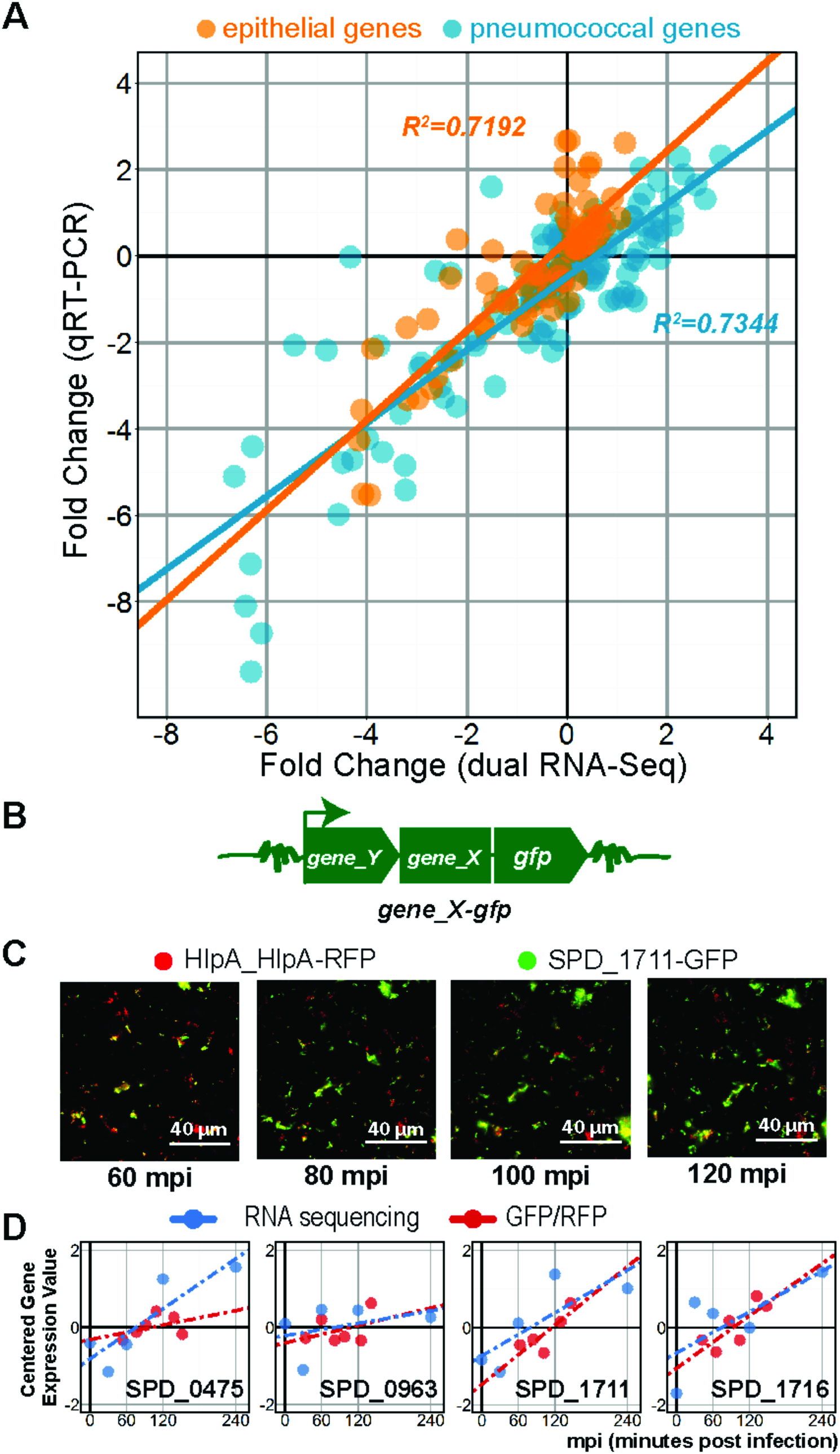
Validation of dual RNA-Seq. **A**. We confirmed dual RNA-Seq gene expression values by qRT-PCR. The infection study was repeated in duplicates and total RNA was isolated as previously. Ten human and 19 pneumococcal genes were chosen as validation targets. We plotted fold changes from qRT-PCR against dual RNA-Seq fold changes and observed a high degree of correlation (*R*^*2*^ > 0.7, Pearson) for both species. **B.** We also confirmed pneumococcal gene expression at the protein level by quantitative fluorescence microscopy. Four target genes (SPD_0475, SPD_0963, SPD_1711 and SPD_1716) were C-terminally tagged with GFP at their own locus. GFP-fusion were performed in the Δ*cps2E* strain expressing RFP fused to HlpA. **C.** Non-deconvolved image of SPD_1711*-*GFP in Δ*cps2E* strain up to 120 mpi. While RFP emitted a relatively constant signal, the GFP signal increases. **D**. We plotted dual RNA-Seq expression values superimposed to the GFP/RFP ratio. To some extent, transcriptional changes corresponded to protein expression.

Since transcript levels do not necessarily correspond with protein expression (Ning et al., 2012; Taniguchi et al., 2010), we quantified four pneumococcal protein levels whose genes showed upregulation during adherence to epithelial cells. We fused a fast-folding variant of the green fluorescent protein (GFP) to the carboxy-termini of SPD_0475, SPD_0963, SPD_1711 and SPD_1716, at their own locus while preserving all upstream regulatory elements (**Figure 3B**). We transformed these constructs into the *hlpA_hlpA-rfp, Δcps2E* genetic background (Kjos et al., 2015). SPD_0475 encodes a 204 amino acids (aa) CAAX amino terminal protease with unknown function. SPD_0963 encodes a 45 aa hypothetical protein. SPD_1711 (132 aa) was described as a single stranded DNA binding protein and may assist in competence (Attaiech et al., 2011) and SPD_1716 is a 63 aa ortholog of cell wall or choline binding protein in other *Streptococcaceae*.

We imaged adherent *S. pneumoniae* with fluorescence microscopy during the indicated time points (**Figure 3C**). Since (i) RFP serves as an accurate proxy for cell number and viability(Kjos et al., 2015), (ii) *hlpA* does not change during early infection (*p*>0.05, *FC*<2) and (iii) ratio between the GFP and RFP indicates relative expression of the protein of interests (red circle and line, **Figure 3D**), we were able to quantify the proteins of interest. Gene expression values from the dual RNA-Seq data (blue circles and line, **Figure 3D**) show a degree of correlation with protein level in three out of the four cases – suggesting that pneumococcal transcriptional changes reflects, to some extent, changes in protein level (Vogel and Marcotte, 2012).

### Pneumococcal ROS induces expression of glutathione-mediated detoxification genes in epithelial cells

Along with pneumococcal adherence and multiplication, we aimed to recapitulate the host response in our model. We hypothesized that the epithelial transcriptome adapts in response to bacterial presence, independent of adherence. To identify the responsive genes, we automatically clustered epithelial working libraries exposed to wild type pneumococci. 242 epithelial genes were co-expressed in a similar manner (**Figure 4A**), i.e., lowly expressed at 30 mpi then sustained upregulation thereafter. Gene ontology (GO) enrichment(Dennis et al., 2003) indicated that 26 of the subset are associated with oxidation and reduction (*p*=5.7·10^−4^). Moreover, 9 genes are associated directly with glutathione, an ubiquitous antioxidant: *GCLC* and *GCLM*, in glutathione biosynthesis; *GPX2* and *MGST2* in detoxification; and *GSR*, *IDH1*, *IDH2*, *PGD* and *G6PD* in glutathione recycling.

**Figure 4.**
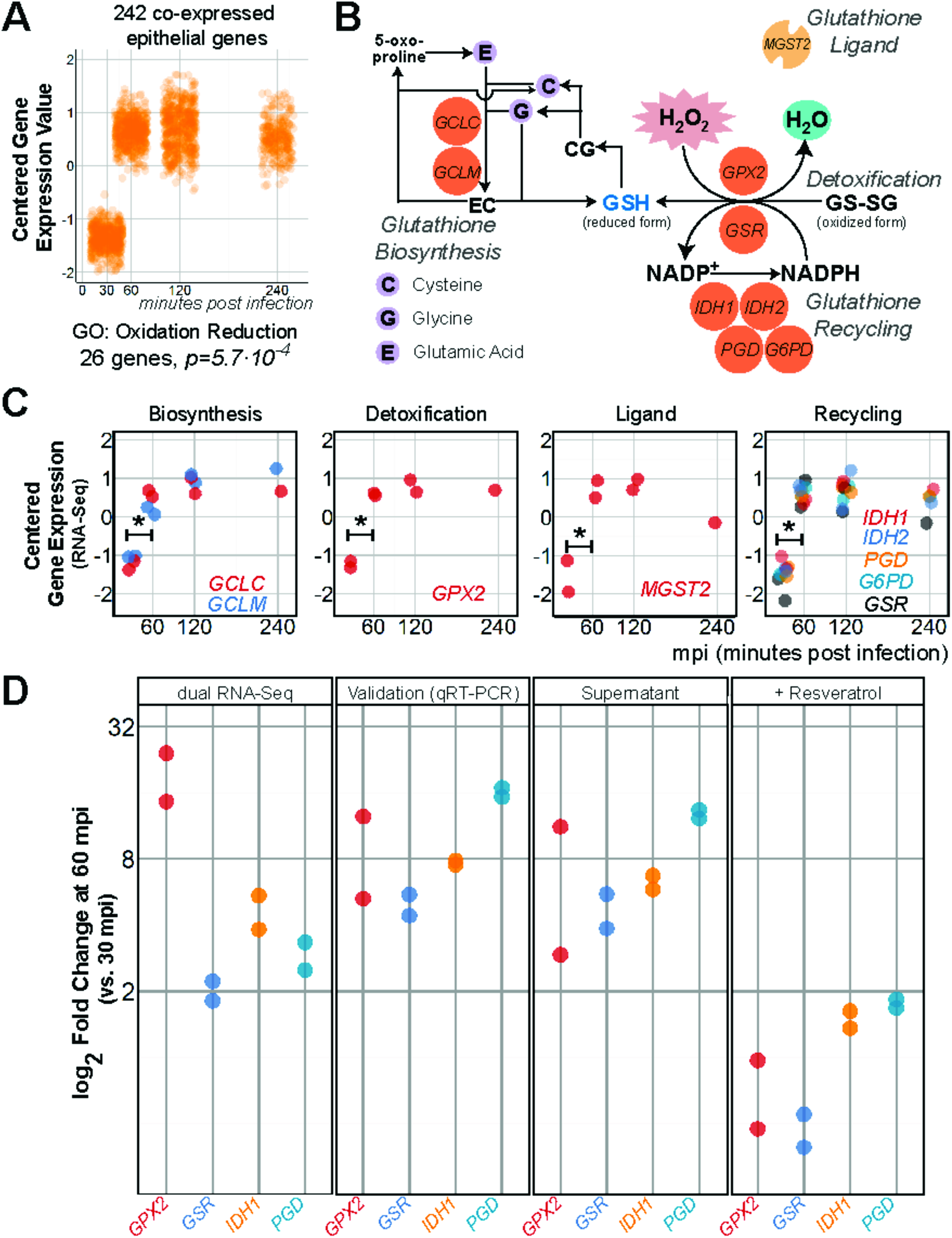
Epithelial glutathione-associated genes are activated in response to pneumococcal ROS. **A**. We clustered epithelial working libraries exposed to wild type pneumococci and found a cluster of 242 co-expressed genes showing sustained upregulation (*p*<0.05, *FC*>2) at 60 mpi compared to 30 mpi. GO analysis showed that “oxidation reduction” was enriched in 26 genes (*p*=5.7·10^−4^). **B**. Nine genes from the subset are associated with glutathione (GSH), an important antioxidant. Main glutathione-associated processes are biosynthesis of glutathione, detoxification of ROS assisted by ligand and glutathione recycling. **C**. Between 30 and 60 mpi, *GCLC* was increased 2.3±1.1 times and *GCLM* 2.4±1.2 times. *GPX2*, the main detoxification gene was activated 18.7±1.3 times while the ligand, *MGST2* 3.2±1.2 times. Genes involved in the recycling of glutathione were activated: *IDH1*, 4.6±1.2; *IDH2*, 2.4±1.2; *PGD*, 2.9±1.2; *G6PD*, 6.6±1.2 and *GSR*, 2.0±1.1 times. **D**. We validated *GPX2*, *GSR, IDH1* and *PGD* expression with qRT-PCR. Epithelial incubation with pneumococcal supernatant showed similar upregulation of glutathione-associated genes. Addition of resveratrol (100μM) into the model diminished the upregulation (*FC*<2) altogether.

Glutathione, a tripeptide of glutamic acid, cysteine and glycine, is produced and secreted by epithelial cells (Valko et al., 2007). The vital molecule is biosynthesized through amino-acid polymerization and when subjected to ROS (hydrogen peroxide, lipid superoxide or oxygen radicals), glutathione readily donates an electron or hydrogen atom to quench the ROS. The process is assisted by ligands and glutathione peroxidase (GPX2). Oxidized glutathione can be recycled by glutathione reductase (GSR) dependent on NADPH (**Figure 4B**). Alternatively, glutathione conjugates and neutralize ROS(Forman et al., 2009). Expression of nine glutathione-associated genes showed a sustained significant increase in epithelial cells exposed to encapsulated strain (*p*<0.05, 60 vs. 30 mpi, **Figure 4C**).

To validate the abovementioned finding, we repeated the experiment, isolated total RNA and performed qRT-PCR on four genes: *GPX2*, involved in detoxification and *GSR*, *IDH1* and *PGD*, in glutathione recycling. As expected, we observed significant upregulation of these genes between 30 and 60 mpi (**Figure 4D**). Interestingly, Rai et al showed that pneumococcal supernatant is sufficient to instigate oxidative damage in A549 (Rai et al., 2015). Indeed, when epithelial cells were incubated with filtered pneumococcal supernatants, the genes were activated (**Figure 4D**). To establish that pneumococci-derived ROS was behind the response, we added the antioxidant, resveratrol (100 μM) to the epithelial-pneumococcal model and did not observe activation (**Figure 4D**) (Zahlten et al., 2015).

### Adherent *S. pneumoniae* repress epithelial innate immune response

Contrasting epithelial gene expression in response to encapsulated and unencapsulated pneumococci allowed us to specifically identify adherence-responsive genes. In doing so, we identified 271 adherence-responsive (*p*<0.05) epithelial genes, of which 25 genes are activated (fold change, *FC>2*) and 248 repressed (*FC>2*) during early-infection (**Figure 5A**). Two genes, *PTGS2* and *HIST1H4B* showed repression and activation at more than one time point. Subsequently, GO-term enrichment in the subset of repressed genes at 60 mpi showed that “immune response” to be enriched in 16 genes (*p*=2.3·10-^4^, **Figure 5B**).

**Figure 5.**
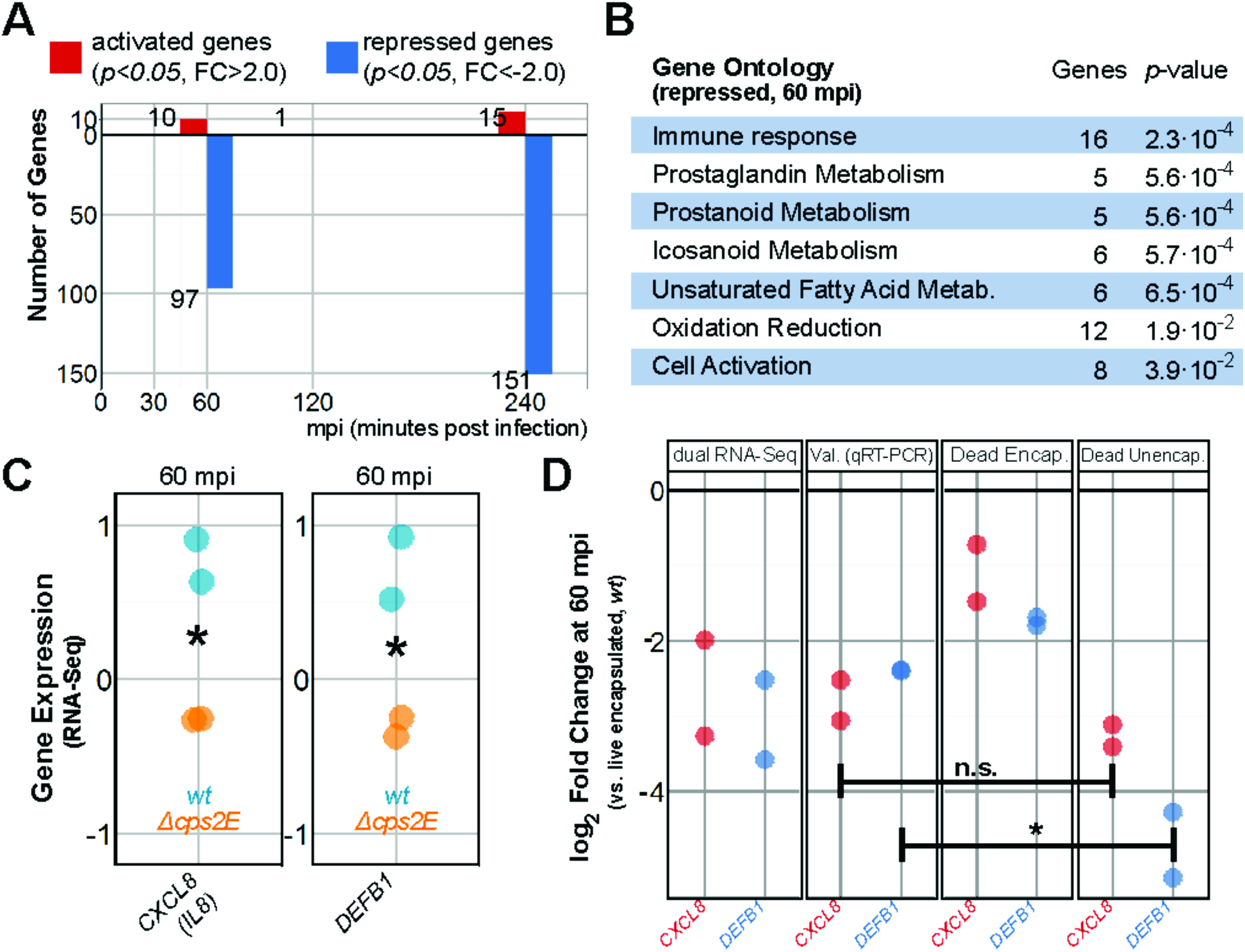
Adherent *S. pneumoniae* repress epithelial innate immune responses. **A**. At 60 mpi, 97 epithelial genes were significantly repressed upon exposure to Δ*cps2E* bacteria compared to exposure to wild type pneumococci. **B.** GO term enrichment to 60 mpi repressed genes resulted in “immune response” (16 genes, *p*=2.3·10^−4^), prostaglandin metabolism (5 genes, *p*=5.6·10^−4^) and oxidation reduction (12 genes, *p*=0.019) among others. **C.** Δ*cps2E-*exposed epithelial cells expressed 2.6±1.3 fold <u>less</u> *CXCL8* and 3.0±1.2 fold <u>less</u> *DEFB1* than wild type-exposed epithelial cells. **D.** We validated *CXCL8* and *DEFB1* repression by qRT-PCR. Heat-inactivated encapsulated bacteria showed no repression of *CXCL8* and *DEFB1*, i.e., no difference (*p*>0.05) compared to viable encapsulated *S. pneumoniae* (Dead Encap.). While infection with heat-inactivated Δ*cps2E* repressed *CXCL8* to the level of viable Δ*cps2E, DEFB1* was more repressed (*p*<0.05) by dead Δ*cps2E* than by viable unencapsulated pneumococci.

*CXCL8 (IL8)*, encoding interleukin-8, was one of the repressed immunity gene. CXCL8 is a potent chemoattractant for neutrophil and other granulocytes. Interestingly, at 60 mpi, Δ*cps2E*-exposed epithelial cells, expressed 2.5±1.3 less *CXCL8* than epithelial cells exposed to wild type *S. pneumoniae* (**Figure 5C**). This difference was validated by qRT-PCR (**Figure 5D**). Further, we asked whether *CXCL8* repression is an active process or merely mediated by physical adherence. To assess this, we co-incubated heat-inactivated Δ*cps2E* and heat-inactivated wild type with epithelial cells. Note that heat inactivation preserves pneumococcal epitope and protein structures (Hvalbye et al., 1999). *CXCL8* was still significantly repressed by dead Δ*cps2E* but not by dead wild type pneumococci (**Figure 5D**) - suggesting that *CXCL8* repression is independent of viability but dependent to presence of the capsule or to the accessibility of surface-exposed (protein) factors in capsule absence. Intriguingly, Graham and Paton showed that epithelial interleukin-8 production and release was suppressed by pneumococcal surface protein CbpA and incubation with Δ*cbpA* leads to higher CXCL8 expression (Graham and Paton, 2006). We speculate that the absence of capsule in Δ*cps2E* increases accessibility of pneumococcal surface-exposed factors, including CbpA to epithelial receptors, leading to repression of *CXCL8*.

*DEFB1*, encoding β-defensin-1 an important epithelial-derived constitutively-expressed antimicrobial peptide (Krisanaprakornkit et al., 1998), is another repressed immunity gene. However, in our model, *DEFB1* was repressed 3.0±1.2 times (*p*<0.05, 60 mpi) in Δ*cps2E*-exposed epithelial cells compared to wild type-exposed cells (**Figure 5C**, validated in **Figure 5D**). Additionally, while heat-inactivated wild type pneumococci stimulate comparable levels of *DEFB1* compared to viable wild type bacteria, non-viable Δ*cps2E* repressed *DEFB1* expression even lower (*p*<0.05) than viable Δ*cps2E* (**Figure 5D**). We conclude that *DEFB1* expression is affected by adherence, accessibility of pneumococcal surface proteins and pneumococcal viability by an as of yet unknown mechanism. In summary, we show that adherent pneumococci modulate epithelial expression of innate immunity genes, including *CXCL8* and *DEFB1*, mediated by pneumococcal surface factors.

### Adherent *S. pneumoniae* activate sugar-importers

Consistent to the epithelial analysis, we contrasted pneumococcal gene expression between unencapsulated and encapsulated libraries, resulting in 248 differentially expressed (*p*<0.05) genes. Specifically, 110 pneumococcal genes of the 248 were activated (*FC*>2) in the unencapsulated strain while 133 genes were repressed (*FC>2*, **Figure 6A**). Five genes, SPD_0188, SPD_0226, SPD_0415, SPD_1717 and SPD_1988, showed repression and activation at different time points. We used gene classes developed for *S. pneumoniae* TIGR4 (van Opijnen and Camilli, 2012) to categorize adherence-responsive genes (**Figure 6B**). Excitingly, most of the adherence-responsive genes are of unknown function (133 of 248 genes, 54%), highlighting our paucity of knowledge in pneumococcal gene function. A large part of the subset (36 genes, 15%) are involved in cellular transport (**Figure 6B**). Since the pneumococcal genome has an exceptionally high number of carbohydrate transporters (Bidossi et al., 2012), it is not surprising that 15 of the genes encode for sugar transporters (**Figure 6C**).

**Figure 6.**
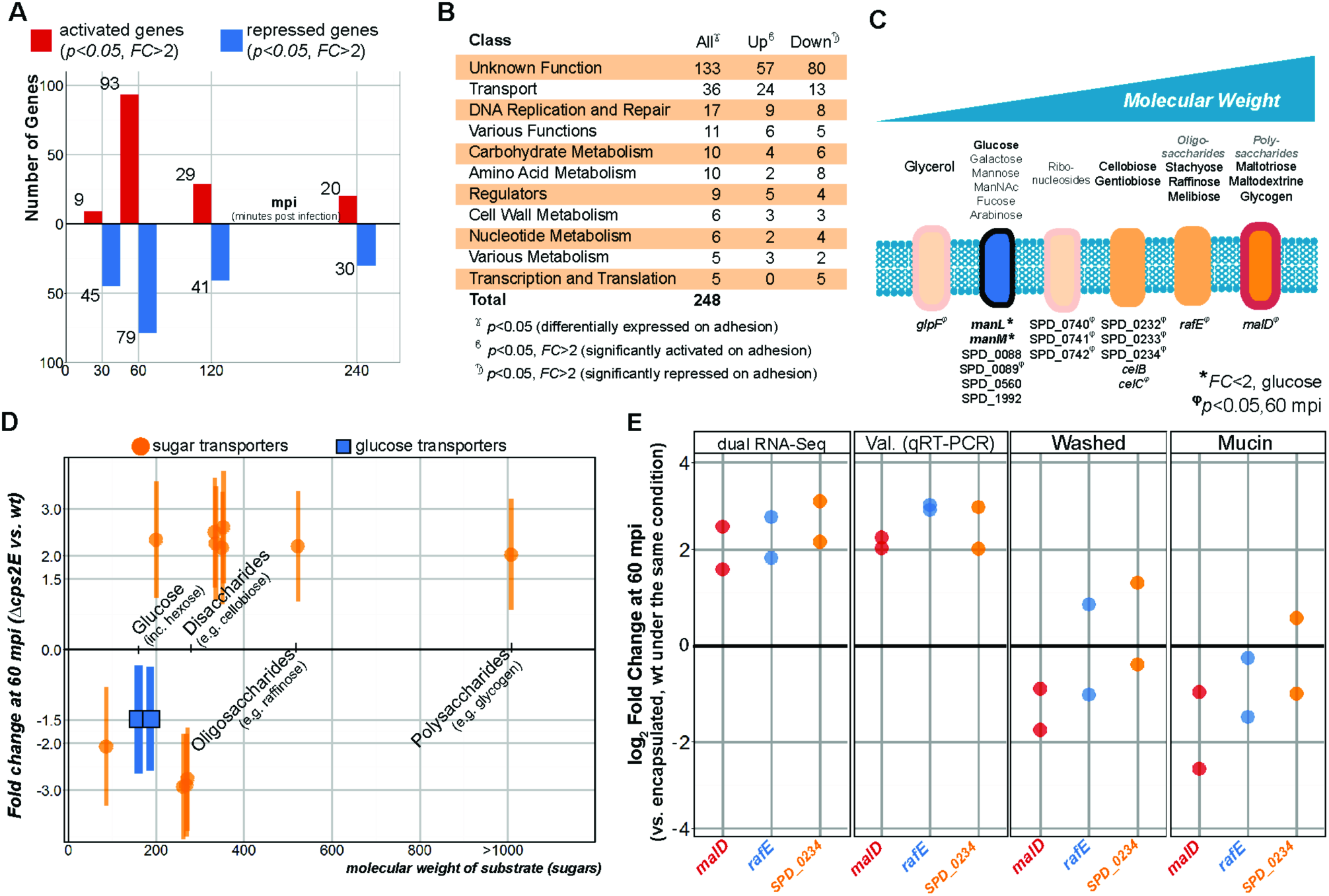
Adherent pneumococci gain access to host-derived carbohydrates and activate non-glucose sugar importers. **A**. 248 genes were differentially expressed between pneumococcal strains exposed to epithelial cells: 115 genes of the 248 genes were activated in Δ*cps2E* compared to wt pneumococci while 138 genes were repressed. Note that five genes showed activation and repression at different time points. **B.** Most of the differentially-expressed genes are of unknown function (133 genes, 54% of 248), followed by cellular transport (36 genes, 15%) and DNA replication, repair and recombination (17 genes, 7%). **C.** 15 of the 36 transporter genes are described to transport carbohydrate. The carbohydrate-importers transport a wide range of carbohydrates, from simple monosaccharides to complex polysaccharides. **D.** At 60 mpi, the expression of glucose transporters (*manLM*, blue boxes) is repressed (*p*<0.05, *FC*=1.5) in Δ*cps2E* compared to encapsulated *S. pneumoniae*. Seven non-glucose transporters are activated (*p*<0.05, *FC>2*) in the Δ*cps2E* strain: SPD_0089, *celC*, SPD_0232/33/34, *rafE* and *malD*. **E.** We validated the data by qRT-PCR for three sugar importers: *malD* (polysaccharides), *rafE* (oligosaccharide), and SPD_0234 (non-glucose disaccharide). By removing epithelial mucus prior to infection, the importers were no longer activated in Δ*cps2E* compared to wild type (*FC*<2, Washed). Incubation with type III porcine mucin (5g·L^−1^) did not activate the genes in Δ*cps2E* compared to encapsulated pneumococci (*FC*<2).

At 60 mpi, 11 carbohydrate transporters were differentially expressed (*p*<0.05, *FC>2*) between Δ*cps2E* and encapsulated pneumococci exposed to epithelial cells. In the presence of high glucose (2g·L^−1^), Δ*cps2E* expressed 1.5 fold less *manLM*, encoding glucose transporters, than encapsulated *S. pneumoniae* (**Figure 6D**). Moreover, seven importers were activated in the adherent strain compared to the free-floating encapsulated strain. The seven non-glucose transporter-genes and their substrates are SPD_0089 (<u>disaccharides</u>: galactose, mannose, N-acetylmannosamine), *celC* (<u>disaccharides</u>: cellobiose, gentiobiose), SPD_0232/33/34 (<u>disaccharides</u>: cellobiose), *rafE* (<u>oligosaccharides</u>: raffinose, stachyose, melliobiose) and *malD* (<u>polysaccharides</u>: maltotitriol, maltodextrine, glycogen, **Figure 6D**). At the same time, four genes were repressed in unencapsulated *S. pneumoniae* exposed to human epithelial cells, *glpF* (glycerol) and SPD_0740/41/42 (ribonucleosides). We selected three transporter-genes, *malD* (polysaccharides), *rafE* (oligosaccharides) and SPD_0234 (disaccharides) and validated the abovementioned observations by qRT-PCR (**Figure 6E**).

Our data indicate that adherent unencapsulated bacteria detect non-glucose sugars in their immediate vicinity. Epithelial mucus may provide non-glucose carbohydrates (Yesilkaya et al., 2008) and simultaneously limit the interaction of epithelial cells to encapsulated wild type bacteria(Nelson et al., 2007). We, then, removed epithelial-associated mucus by washing the surface with warm PBS and observed that the genes were no longer activated (*FC*<2, **Figure 6E**). Next, we incubated pneumococcal strains with type III porcine mucin (5 g·L^−1^), mimicking complex carbohydrate in the medium. Interestingly, the importers were not differentially expressed between strains (*FC*<2), indicating similar access to non-glucose sugars (**Figure 6E**). We conclude that following adherence, *S. pneumoniae* senses an enriched host-derived non-glucose carbohydrate and in turn, activates transporters to import the now-available sugars.

## Discussion

Early infection is a chaotic and disruptive encounter between host and pathogen. In both species, a multitude of transcriptional-mediated cellular processes are fine-tuned: activated, maintained and repressed to ensure survival. The recently-described dual RNA-Seq approach allows simultaneous host-pathogen monitoring during their interactions (Baddal et al., 2015; Dillon et al., 2015; Westermann et al., 2016). In this study, we exploited the dual RNA-Seq approach by application to a pneumococcal infection model to human lung alveolar epithelial cells. We have generated a detailed time-resolved dataset of epithelial-pneumococcal transcriptomes up to 4 hours after infection. Moreover, we have validated the rich dataset by qRT-PCR and quantitative fluorescence microscopy. Furthermore, we have shown that adherence-specific transcriptional responses in host and pathogen can be identified by contrasting the transcriptomes of the encapsulated libraries with its unencapsulated counterpart.

Our early-infection model recapitulated three major *in vivo* characteristics of pneumococcal infection: pneumococcal adhesion, bacterial multiplication and epithelial responses to pathogenic burden. Our model recapitulated these infection characteristics: (i) adherence for both encapsulated and unencapsulated pneumococci (**Figure 1**), (ii) pneumococcal viability and growth during early infection, e.g.: generation of ROS, expression of carbohydrate importers (**Figures 4** and **6**) and (iii) host response to pneumococcal presence, e.g.: glutathione-associated detoxification and innate immune response (**Figures 4** and **5**). Further, as shown by regulation of carbohydrate transporters (**Figure 6**), pneumococci sensed epithelial presence and subsequently adapted their transcriptome.

Remarkably, we observed almost all (99%) pneumococcal genes are expressed at early infection (**Figure 2**) – in line with recent studies on bacterial transcriptomes adapting to multiple conditions(Kröger et al., 2013; Nicolas et al., 2012). We speculate that interspecies interaction necessitates massive pneumococcal transcriptional adaptation. Moreover, we have observed activation of detoxification genes (*GPX2* and *GSR*) in epithelial cells protecting against pneumococci-derived ROS (**Figure 4**). Indeed, *S. pneumoniae* has been reported to secrete high levels of peroxides as a by-product of its pyruvate metabolism (Carvalho et al., 2011) and has recently been shown to cause DNA-damage-dependent apoptosis in alveolar lung epithelial cells(Rai et al., 2015). In addition, pneumococcal early competence genes (Martin et al., 2000) were activated in our model (http://dualrnaseq.molgenrug.nl competence subset). Future work should also examine whether small non-coding RNAs play a role in pneumococcal early infection as they do in Salmonella (Westermann et al., 2016), something that was not explored currently due to the poor D39 genome annotation in this regard.

Our approach can be expanded further by incorporating relevant LRTI agents into the model. For example, alveolar macrophages and epithelial cells, together, form epithelium lining of the lower respiratory tract. The cells reciprocally influence cellular phenotypes and behaviors (Hussell and Bell, 2014), highly relevant to infection. Moreover, pneumococcal co-infection and secondary infection are not unheard of. Influenzae virus potentiates *S. pneumoniae* by neuraminidase activity that exposes cryptic epithelial receptor and facilitates pneumococcal adherence(Siegel et al., 2014). In addition, influenzae infection causes loss of superficial epithelial cells, revealing basal epithelium (Kash et al., 2011) and aggravating secondary pneumococcal infection. On the other hand, *H. influenzae* negatively affects pneumococci by recruiting neutrophils and stimulating the killing of opsonized pneumococci(Lysenko et al., 2010). Incorporation of other agents into the model and exploiting dual (or triple, quadruple) RNA-Seq approaches may provide novel insights into respiratory infection.

However, complementary approaches, are needed to obtain a full picture of early infection. Though transcriptome rewiring is a focal point during interspecies interaction (Jenner and Young, 2005; Sorek and Cossart, 2010), non-transcriptional regulation plays an important part during early infection. Capsule shedding, a hallmark of pneumococcal infection, is regulated by autolysin-A (Lyt-A). LytA is activated when the bacterium encounters alveolar cathelicidin (Kietzman et al., 2016), which is independent of transcriptional regulation. Heterogeneity of cellular responses is another confounding factor (Jørgensen et al., 2013). Recently, dual RNA-Seq combined with cell sorting was used to identify heterogeneous activity of Salmonella virulence factor that, in turn, drives heterogeneous interferons response in macrophage(Avraham et al., 2015). This highlights the relevance of noise in gene expression and cell-to-cell variability in host-pathogen interaction. Furthermore, whole organism infection models offers a more systemic perspective. Dual RNA-Seq approach has been used to monitor infection in wheat-bacteria (Camilios-Neto et al., 2014) and mosquito-filaria (Choi et al., 2014). Whole organism dual RNA-Seq is not without its challenges, including averaging (host) effects to gene expression across all cell types.

Besides its relevance in communicable diseases, gained insights into pneumococcal infection are also applicable to understanding several non-communicable respiratory diseases. Asthma, the most common chronic respiratory disease is a major risk factor for pneumococcal infection (Talbot et al., 2005). Additionally, COPD (Chronic Obstructive Pulmonary Disease, Decramer et al., 2012) and cigarette smoking (Phipps et al., 2010) have been reported to increase the risk of pneumococcal LRTI. Here, we reported the first study to show simultaneous transcriptomic changes of the pathogen *S. pneumoniae* and human lung alveolar epithelial cells during early infection. Additionally, we have made the time-resolved dual transcriptomics dataset available to the broader research community (http://dualrnaseq.molgenrug.nl). We invite pneumococcal researchers to use the database to formulate research questions in the development of preventive and curative strategies against pneumococcal infection. To conclude, we call researchers from the fields of microbiology, immunology and pulmonology to access the dataset and use it to develop their own hypotheses.

## Methods

### Culture of epithelial cell line, *S. pneumoniae* D39 and pneumococcal transformation

Human type II lung epithelial cell line, A549 (ATCC^®^ CCL-185) and *S. pneumoniae* D39 were routinely cultured without antibiotics. Strain construction is described in detail in **Supplemental information**. Oligonucleotides are listed in **Supplemental Table 1** and strains in **Supplemental Table 2**.

### Infection studies

Confluent monolayer of A549 was co-incubated with *S. pneumoniae* D39 at multiplicities of infection 10, in 1% fetal bovine serum in RPMI1640 without phenol red. Prior to infection, epithelial monolayer was kept for 10 more days after confluence. To optimize cell-to-cell contact, centrifugation was employed (2000 ×g, 5 min, 4°C). Adherence assays were performed by enumeration of plated cfu in blood agar. More in **Supplemental Information**.

### Simultaneous total host-pathogen RNA isolation

Before RNA isolation, samples were treated by saturated ammonium sulfate solution (Korfhage et al., 2002). Cells were disrupted by bead beating, and total RNA was isolated as described in the **Supplemental Information**.

### Library preparation, sequencing, data pipeline and online database

Total RNA was dual rRNA-depleted, reverse-transcribed and sequenced on the Illumina NextSeq 500 in 75 nt single end mode. Raw reads were trimmed, aligned into chimeric human-pneumococcus genome. Reads were counted and differential gene expression analysis were performed to epithelial and pneumococcal libraries separately. Automatic clustering and GO enrichment were performed. The online database is built using in-house script. Significantly differentially expressed epithelial and pneumococcal genes are listed in **Supplemental Table 3** and **4**, respectively.

### qRT-PCR and quantitative fluorescence microscopy

Infection studies were repeated, total RNA isolated and qRT-PCR is performed. For fluorescence microscopy, infection studies were performed in 8-wells μ-slides (Ibidi, DE). More in **Supplemental Information**.

## Data Access

The raw data is accessible at http://www.ncbi.nlm.nih.gov/geo/ with accession number GSE79595.

## Author Contribution

R.A and J.W.V. designed the research, analyzed the data and wrote article. R.A performed research, J.S analyzed the data, S.H. built online database.

## Acknowledgements

We thank W.J. Quax and R. Setroikromo (UMCG, Groningen) for the human cell line, V. Benes and B. Haase (GeneCore, EMBL, Heidelberg) for sequencing support, M. Kjos for fruitful discussions and A. de Jong for bioinformatics support. We thank S. El Aidy and L.E. Keller for comments on the manuscript. Work in Veening lab is supported by the EMBO Young Investigator Program, VIDI fellowship (864.12.001) from the Netherlands Organization for Scientific Research, Earth and Life Sciences (NWO-ALW), and ERC Starting Grant 337399-PneumoCell.

## Conflict of Interest

None

